# *Pseudomonas* two-partner secretion toxin Exolysin contributes to insect killing

**DOI:** 10.1101/807867

**Authors:** Viviana Job, Stéphanie Bouillot, Erwan Gueguen, Mylène Robert-Genthon, Peter Panchev, Sylvie Elsen, Ina Attrée

## Abstract

*Pseudomonas chlororaphis* is a promising biocontrol agent promoting plant-growth and providing protection against pest insects and phytopathogenic fungi. We have identified in the genome of *P. chlororaphis* PA23 an operon encoding the toxin Exolysin (ExlA) and its outer-membrane transporter, ExlB. We found that *P. chlororaphis* producing ExlA (ExlA^*Pch*^) is cytotoxic towards murine macrophages and human epithelial cells at 30 °C. *P. chlororaphis* PA23 provoked shrinkage of epithelial cell, leakage of cytoplasmic components and subsequent cell death. During infection, ExlA^*Pch*^ incorporated into epithelial cell membranes within detergent-resistant lipid rafts, suggesting the same mechanisms of cell destruction by pore-formation as reported for *P. aeruginosa* toxin. ExlA^*Pch*^ was not involved in the capacity of the strain to kill fungi, amoeba or other bacteria. The contribution of ExlA in insecticidal activity of *P. chlororaphis* was evaluated in the wax moth larvae *Galleria mallonella* and in *Drosophila melanogaster* flies. The impact of the deletion of a gene encoding *exlA* homologue was tested in the natural fly pathogen *P. entonomophila*. In both models, the ExlA absence delayed killing, suggesting the contribution of the toxin in bacteria-insect pathogenic interactions.

## Introduction

Bacteria from the genus *Pseudomonas* are ubiquitous in nature due to their extraordinary capacities to adapt to diverse niches. *Pseudomonas* species are mainly associated with plants and animals but also frequently found in the proximity of human activities [1]. *P. fluorescens, P. putida* and *P. stutzeri* are recognized as beneficial with potential uses in bioremediation, production of secondary metabolites, or in agriculture for the plant-growth promoting activity and biocontrol property (reviewed in [1–4]). To the contrary, *P. aeruginosa* is a well-known human opportunistic pathogen that causes high-cost health problems notably due to the increasing number of multi-drug resistant strains. The World Health Organization [WHO] classified carbapenem-resistant *P. aeruginosa* in a high priority category of bacterial pathogens for which there is an urgent need to define new antibiotics [5].

*P. chlororaphis* belonging to the group complex of *P. fluorescens* thrives on plants and in diverse soil environments. Several strains of *P. chlororaphis* have been selected as potential biocontrol agents as they possess both pest insecticidal activities and are able to suppress the plant fungal pathogen *Sclerotinia sclerotiorum* [6–8] that causes the disease called “white mold” in a wide range of plants. The spectrum of secondary metabolites produced by *P. chlororaphis* includes phenazines, pyrrolnitrin, hydrogen cyanide and others [7, 9, 10]. In addition, secreted proteinous macromolecules as chitinases, proteases and toxins participate to *P. chlororaphis* success in phytoprotection and promotion of plant growth [9].

Together with phenotypic studies, the accessibility to several fully sequenced *P. chlororaphis* strains available at Pseudomonas database, www.pseudomonas.com [11], allowed the analysis of their genome content with respect to predicted secreted molecules and toxins. The reference strain *P. chlororaphis* PA23 [12] harbors the gene cluster encoding a toxin, Fit, and the predicted dedicated export machinery identified originally in *P. protegens* [13]. The Fit toxin is responsible for killing of several important lepidopteran pests, notably *Spodoptera littoralis* (cotton leafworm) and the diamondback moth *Plutella xylostella* [14]. *P. chlororaphis* PA23 genome also harbors some genes potentially encoding type III secretion system (T3SS) components and two loci encoding T6SS. In *P. chlororaphis*, T6SSs participate to the invasion of a Gram-positive bacteria *Bacillus subtilis* [15] and may play a role in displacement of the pest insect gut microbiome as recently described for *P. protegens* [16]. Moreover, high-molecular weight bacteriocins, named Tailocins, could broaden the killing capacities of *P. chlororaphis* in polymicrobial communities [17, 18].

Clinical strains of the human opportunistic pathogen *P. aeruginosa* use either T3SS or two-partner secretion (TPS) system ExlB-ExlA to promote host invasion and infection [19, 20]. The presence of *exlBA* operon in *P. chlororaphis* suggested its role in establishing pathogenic relationship with other organisms, prokaryotes or eukaryotes. In this work, we examined the expression and functionality of ExlA from *P. chlororaphis* and showed its contribution in killing of *Galleria mellonella* larvae upon injection. Moreover, ExlA homologue in *P. entonomophila*, a fly pathogen [21] was found to play a role in oral toxicity of *Drosophila melanogaster* flies.

## Results and discussion

### *P. chlororaphis EY04_RS20945* and *EY04_RS20950* encode ExlB-ExlA

The search in databases for the closest homologues of *P. aeruginosa* ExlB (ExlB^*Pa*^) and ExlA (ExlA^*Pa*^) revealed two contiguous genes, *EY04_RS20945* and *EY04_RS20950* in *P. chlororaphis* PA23 sharing homologies with *exlBA* in several clonal *P. aeruginosa* outliers (www.pseudomonas.com, [11], Figure 1). *EY04_RS20945* encodes a 65-kDa outer membrane protein homologue to ExlB, a TpsB partner involved in the export of the cognate TpsA secreted protein. *EY04_RS20950* encodes the predicted 170-kDa TpsA secreted partner with a classical Sec secretion signal peptide at the N-terminus followed by the TPS secretion domain, and six predicted FHA repeats. The predicted *EY04_RS20945* and *EY04_RS20950* gene products share 67 % and 62 % identity to ExlB^*Pa*^ and ExlA^*Pa*^, respectively. Taking into account the high identity scores and the data presented in this work, we renamed *EY04_RS20945* and *EY04_RS20950* as *exlB^Pch^* and *exlA^Pch^*, respectively. Genomic environment of *exlB^Pch^* and *exlA^Pch^* is different from that found in *P. aeruginosa* strains, with several regulatory proteins encoded upstream and downstream of both operons (Figure 1). Notably, two genes upstream of *exlB^Pch^* encode enzymes involved in biogenesis of the second messenger c-di-GMP, while two genes downstream of *exlA^Pch^* encode putative two-component regulator system with a histidine kinase (HK) and a transcriptional factor (TF). The region upstream of *exlB^Pch^* ATG start codon harbors a putative promoter (BPROM) with predicted “-35” and “-10” σ^70^-RNA polymerase binding sites. The *exlBA* locus is present in all *P. chlororaphis* strains reported at the *Pseudomonas* genome database and the syntheny of the locus is shared by *P. protegens* strains (www.pseudomonas.com). We also reanalyzed the region of two genes *PSEEN_2176* and *PSEEN_2177* in a fly pathogen *P. entonomophila* that code for a two-partner system homologue to ExlB-ExlA ([22], Figure 1). PSEEN_2176 gene product shares 69% identity with ExlB^*Pch*^ and 65 % with ExlB^*Pa*^, while PSEEN_2177 codes for an ExlA like-protein with 47 % and 40 % identity with ExlA^*Pch*^ and ExlA^*Pa*^, respectively. We renamed also these genes product as ExlA^*Pe*^ and ExlB^*Pe*^.

**Figure 1.**
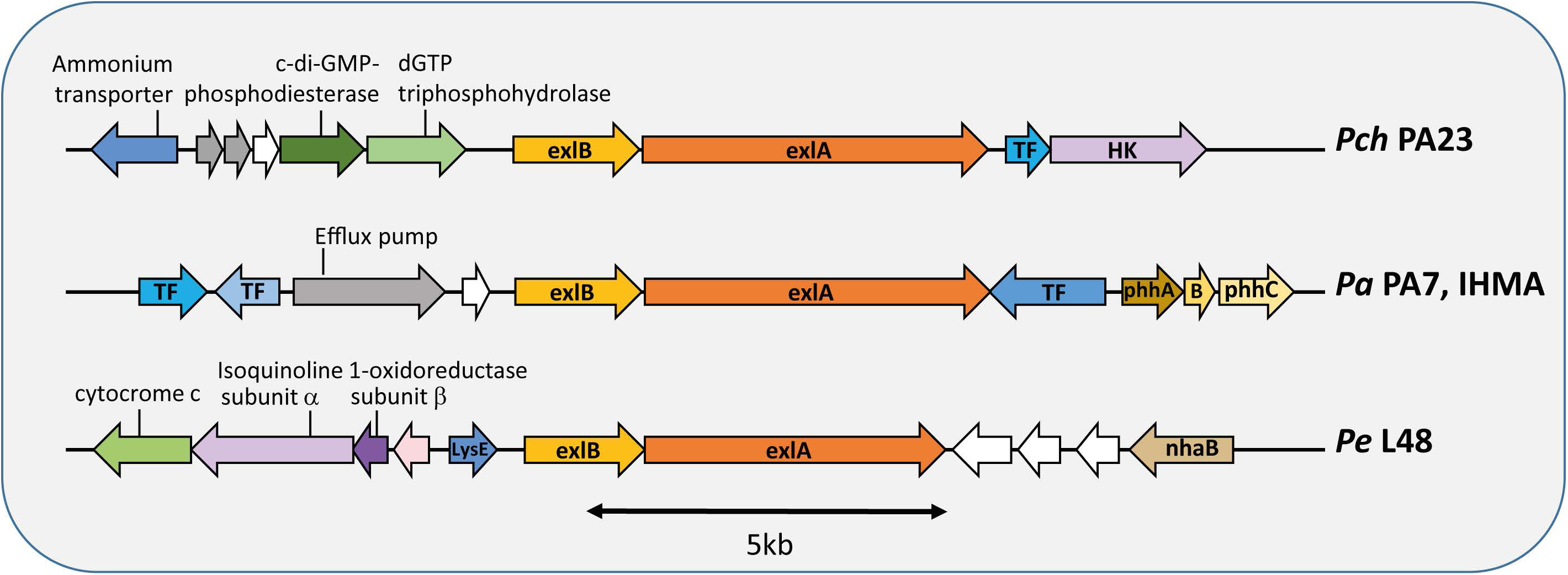
Genetic environment of the *exlB*-*exlA* operon. The sequences corresponding to *P. chlororaphis* PA23 (*Pch*), *Pseudomonas aeruginosa* PA7 (*Pa*) and *Pseudomonas entomophila* L48 (*Pe*) were retrieved from *Pseudomonas* genome database (www.pseudomonas.com) and annotated accordingly. The conserved genes *exlA* and *exlB* are in dark and light orange, respectively. Blue color indicates predicted putative regulators. Genes of unknown functions are in grey and encoding hypothetical proteins in white. Scale is shown.

### ExlA provides cytotoxic phenotype to *P. chlororaphis* PA23

Virulent *P. aeruginosa* clinical isolates secreting ExlA (ExlA^*Pa*^) display cytotoxicity on a variety of eukaryotic cells by forming pores in the host plasma membrane leading to altered Ca^2+^ signaling, cleavage of cell-cell junctions and cell death [23, 24]. To assess the cytotoxic potential of *P. chlororaphis* on eukaryotic cells, we co-incubated at 30 °C epithelial A549 cells with *P. chlororaphis* PA23 ranging from multiplicity of infection (MOI) of 0.1 to 100, and measured the kinetics of incorporation of the membrane-impermeable, propidium iodide (PI) into DNA (Figure 2A). PI incorporation was detected already with MOI of 0.1, meaning 1 bacteria per 10 epithelial cells, and the kinetics progressively increased with higher MOI. To ascertain that the observed cytotoxicity of *P. chlororaphis* PA23 was due to the ExlA protein, we engineered a strain in which a part of the *exlB*-*exlA* operon was deleted (*Pch*Δ*exlBA*) and measured the kinetics of PI incorporation during 3 h in comparison to *P. aeruginosa* strain IHMA87 and the *exlA*-isogenic mutant (*Pa*-*exlA*mut, [23, 25]) on macrophages and epithelial cells. The *Pch*Δ*exlBA* strain lost the capacity to kill J774 macrophages and A549 epithelial cells (Figure 2B). We then followed the infection process by video-microscopy. Similar to the time-lapse consequences provoked by the ExlA-secreting *P. aeruginosa*, *P. chlororaphis* induced initial epithelial cell rounding, followed by the loss of cytoplasmic content, which can be visualized by the loss of cytoplasmic GFP. The incorporation of PI into the cell nuclei occurred at the later stages of infection process (Figure 3). As ExlA^*Pa*^ is a membrane pore-forming toxin and binds to liposomes and host membranes ([23], V. Job, manuscript in preparation), we tested whether ExlA^*Pch*^ could be detected within infected cell membranes, and in particular within the lipid rafts, detergent-resistant membrane (DRM) fractions enriched in cholesterol and sphingolipids, that is involved in protein sorting, signaling and interaction with the cytoskeleton. To that end, after scaled-up infection conditions, we prepared and analyzed e epithelial cell membranes for presence of ExlA^*Pch*^ by immunoblotting. Both ExlA^*Pch*^ and ExlA^*Pa*^ could be recovered in DRM fractions (Figure 4), indicating the same mechanism of interaction with cell membranes and similar action of those two proteins with host membranes.

**Figure 2.**
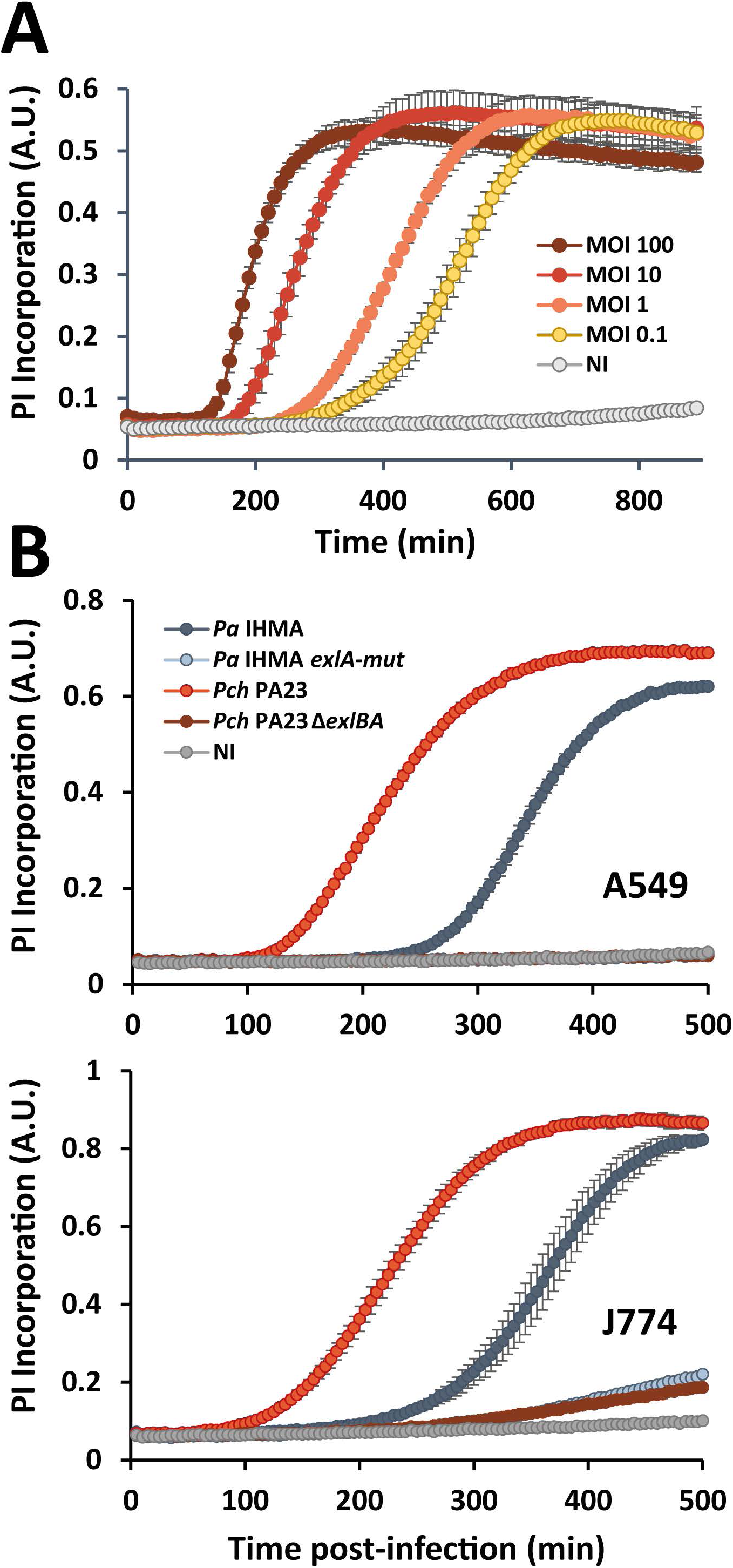
Cytotoxicity of *P. chlororaphis* PA23 on eukaryotic cells is ExlA-dependent. **A.** Kinetics of A549 epithelial cells cytotoxicity depends on MOI. Epithelial cells were infected with increasing quantities of bacteria ranging from MOI of 0.1 to 100. **B.** *P. chlororaphis* is cytotoxic *in vitro* in an ExlBA-dependent manner. Epithelial cells (A549, upper panel) or murine macrophages (J774, lower panel) were infected at a MOI of 10 at 30 °C with *P. chlororaphis* and *P. aeruginosa* strains, as indicated. Cell death was monitored by PI incorporation. Fluorescence emission at 590 nm of PI expressed as arbitrary units (A.U.) was recorded in function of time. Experiments were carried out in triplicates, and the error bars indicate the standard deviation.

**Figure 3.**
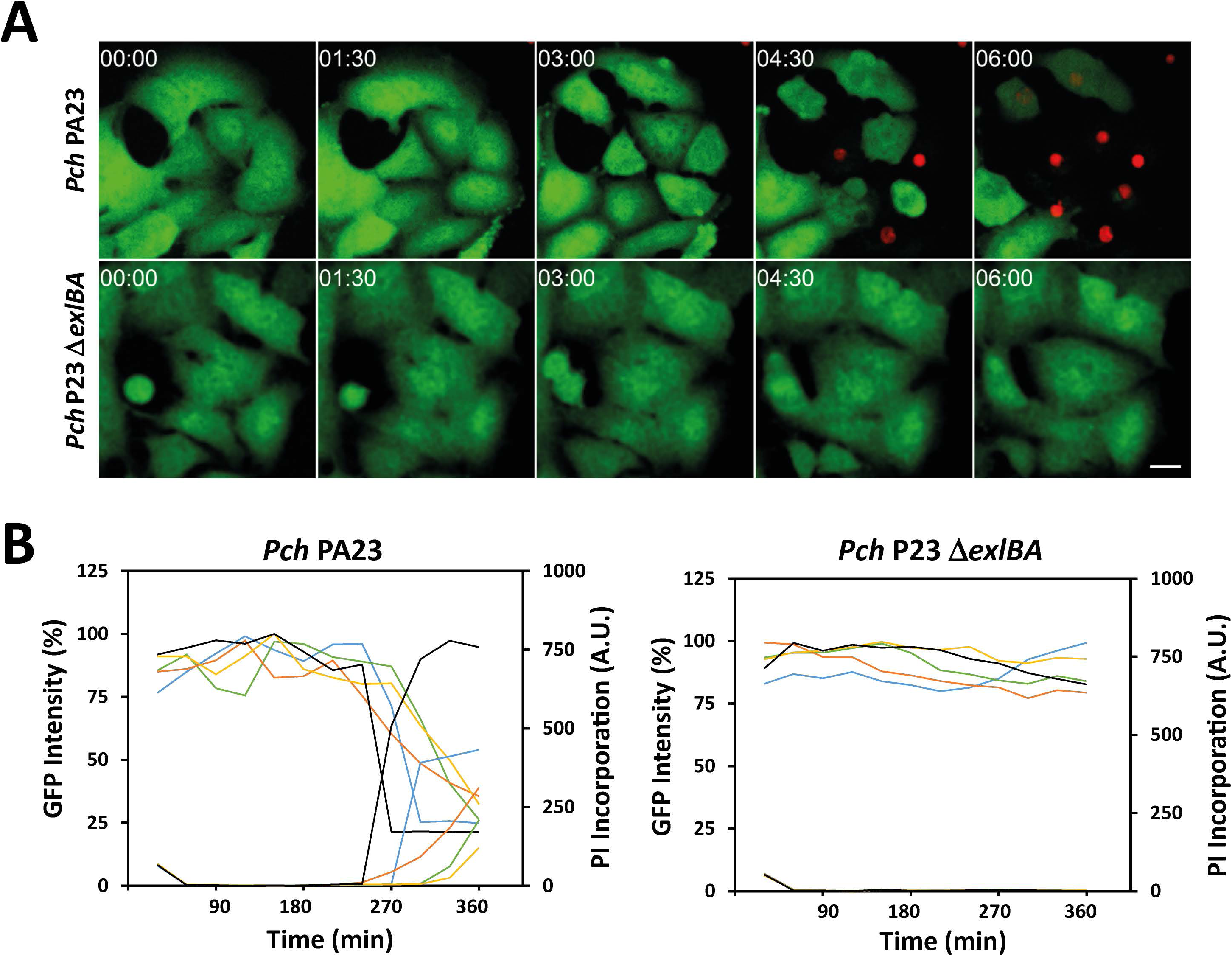
Consequences of interactions between epithelial cells and *P. chlororaphis* PA23 or *exlBA* isogenic mutant. **A**. Epithelial cells A549 expressing cytoplasmic GFP were incubated with *P. chlororaphis* PA23 or *exlBA* mutant at MOI of 10. Incorporation of PI (red) into cell nuclei was followed by confocal videomicroscopy. Times post-infection are shown as “h:min”. One z-position is represented. Scale bar shown on the last image corresponds to 20 µm. Images are representative of five positions simultaneously recorded for each condition. **B.** Intensities of GFP and PI were quantified on time lapse films reported in A. Five cells were analyzed in each condition. The GFP intensities are represented by solid lines and PI intensities by dashed lines, using the same color code for each selected cell.

**Figure 4.**
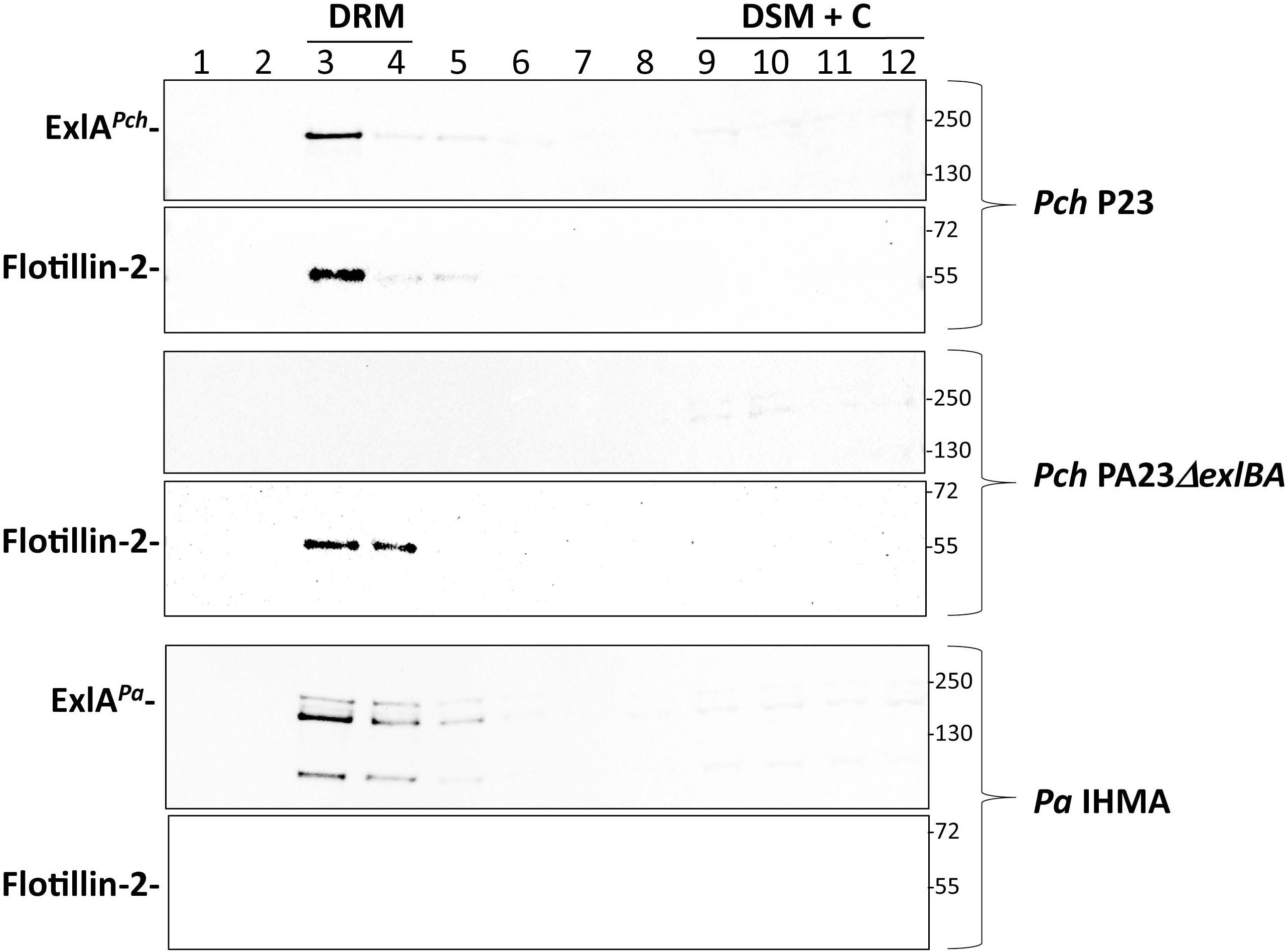
ExlA^*Pch*^ fractionates with lipid raft portion of infected epithelial cell membranes. A549 epithelial cells were infected with *P. chlororaphis* (wild-type or *exlBA* mutant) or *P. aeruginosa* IHMA87 wild-type strain, used as positive control at OD_600_ of 1.0 with an MOI of 10. The fractions with detergent-resistant membranes (DRM) containing lipid rafts, the detergent-soluble membranes (DSM) and the cytoplasm (C) were separated on sucrose gradient after treatment with 1 % Triton X-100, as described in Material and Methods, and analyzed by Western blot using anti-ExlA antibodies mixture. Anti Flotillin-2 antibodies were used to localize DRM-containing fractions. Numbers on the top refers to the twelve fractions recovered from the top to the bottom of sucrose gradient, while sizes of proteins marker in (kDa) are shown on the right of the Western Blot.

### ExlA^*Pch*^ in microbial competition and amoeba killing

*P. chlororaphis* is considered as a safe biocontrol agent [26, 27] and until recently has been rarely found responsible for human infections [28]. *P. chlororaphis* is frequently associated with plants, to which it provides protection against diverse phytopathogens, including fungi and insects [2, 29]. We thus wondered whether ExlA^*Pch*^ could participate in interactions with other microbes. First, we tested an inhibition of *P. chlororaphis* on the plant fungal pathogens *Sclerotinia sclerotiorum* and *Botrytis cinerea* [6, 8], using radial inhibition assay on agar PDA plates (Figure 5A). No significant difference in fungus growth inhibition could be observed between PA23 wild-type strain and the *exlBA* mutant, suggesting that the toxin does not participate in fungicide activity of PA23 under condition tested. We tested also putative implication of ExlA in bacteria-bacteria interactions, using *B. subtilis*, *P. aeruginosa* and *E. coli*. In all three cases, no difference could be detected between the *P. chlororaphis* wild-type and mutant strains.

**Figure 5.**
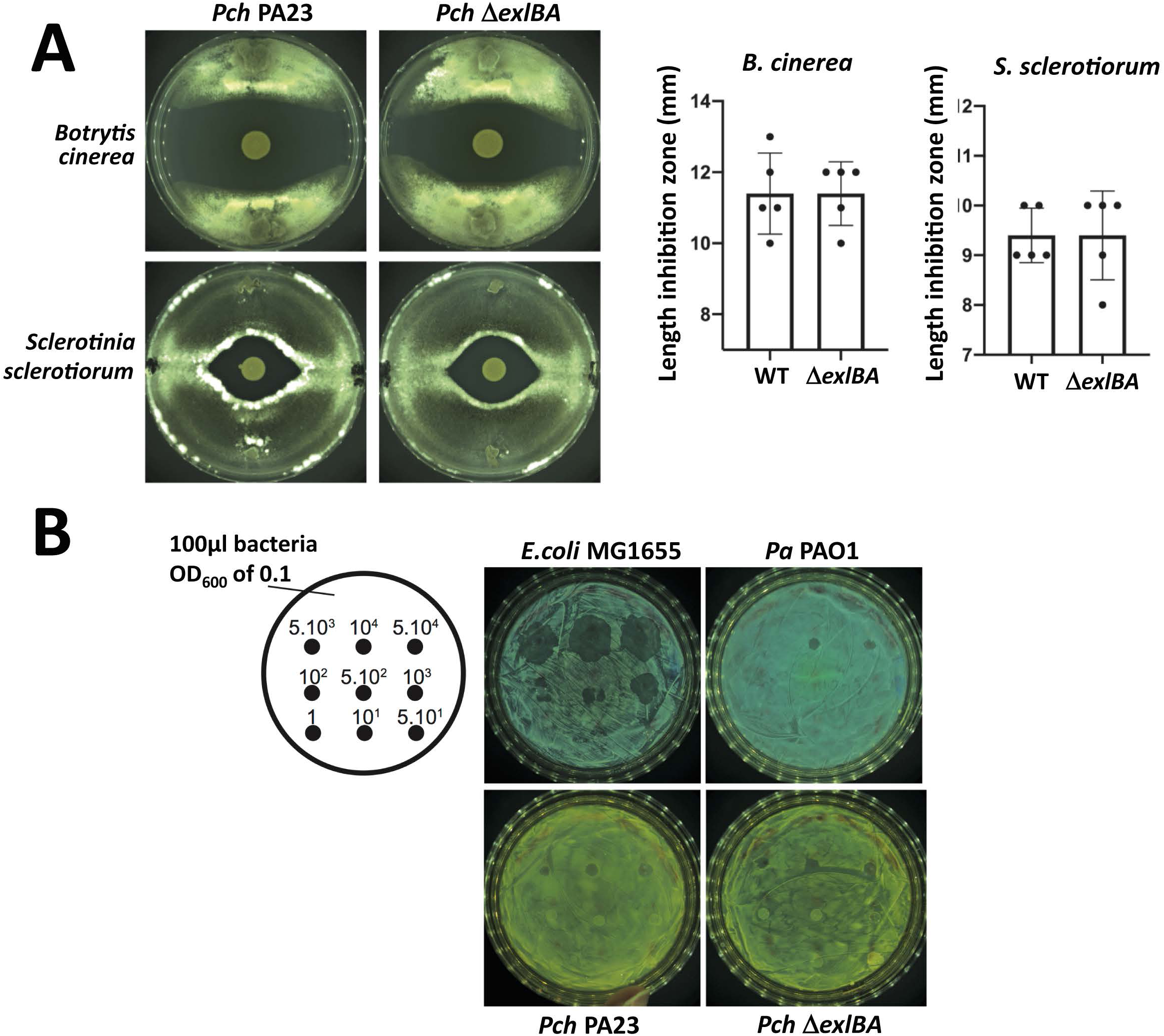
ExlA is not involved in interaction of *P. chlororaphis* PA23 with fungus *Sclerotinia sclerotiorum* and *Botrytis cinerea* nor in resistance to grazing by *Acanthamoeba castellanii*. Radial diffusion assays to assess *Sclerotinia sclerotiorum* inhibition of growth *in vitro* on PDA plates with *P. chlororaphis* wild-type or ∆*exlBA* strains. 5 µl of *P. chlororaphis* at OD_600nm_ of 0.1 was deposited to the center of a 90 mm Petri dish inoculated with either plugs of *Sclerotinia sclerotiorum* or *Botrytis cinerea* mycelium. Plates were incubated at 25 °C until the mycelium covers the plate. Lengths of fungal inhibition zone was measured for 5 independent experiments. No statistical difference was measured between the strains (Mann-Whitney test). **B.** *Acanthamoeba castellanii* plaque formation assay. The plaques formed by amoebae on the lawns of *P. chlororaphis* wild-type or ∆*exlBA* strains. *P. aeruginosa* is used as a negative control since this bacterium kill *Acanthamoeba*. *E. coli* is used as a positive control since this bacterium can be phagocyted. The number of amoebae cells deposited on the plate is indicated (from 5.10^4^ on the top left to 5.10^1^ on the bottom right corner). 100 µl of bacterial culture at OD_600_ of 0.1 was spread onto each plate containing M63 nutrient agar with glucose 0.2 %.

Second, we wondered if ExlA could influence the growth of eukaryotic organisms such as amoeba. For example, *P. aeruginosa* PAO1 is capable of killing the amoeba *Acanthamoeba castellani* with its T3SS [30]. To test whether *P. chlororaphis* could also infect *A. castellani*, the pathogenicity of *P. chlororaphis* toward *Acanthamoeba castellani* cells was determined by using a simple plating assay [31]. As shown in Fig. 5B, 50 amoebae cells are required to form a plaque on *E. coli*, while 10,000 are necessary to form a plaque on *P. aeruginosa* PAO1. This result is explained by the ability *of P. aeruginosa* to kill amoebas by injecting T3SS cytotoxins [31] while *E. coli* are grazed. With *P. chlororaphis*, no difference in plaque formation was observed between wild type and ∆*exlBA*. In both cases, 5,000 amoebas were required to form a small plaque. This result is similar to that obtained with *P. aeruginosa*. This experiment does not distinguish between killing and growth inhibition of *A. castellani*. Nevertheless, it suggests that ExlA does not play a role in the interaction with the amoeba in conditions tested. Of note, the interactions between the *Pseudomonas* and amoeba were done on solid medium and ExlA may not be expressed and/or secreted in this condition. Otherwise, the composition of the host membranes between permissible and not permissible cells may be of crucial importance for ExlA function.

### ExlA contributes to toxicity towards *Galleria mellonella* larvae and *Drosophila* flies

*P. chlororaphis* strains possess oral and systemic insecticidal activity [9]. To assess the contribution of Exolysin to insect killing, we used *G. mellonella* larvae and *Drosophila melanogaster* flies. When *P. chlororaphis* PA23 bacteria were injected into hemocoel of the larvae, the mortality could be observed at 18-20 h post-injection (Figure 6) with typical change in color of the larvae which is directly related to melanization, a larvae’s immune response (Figure 6A). The mortality rates were dependent on temperature and number of injected bacteria (data not shown). The absence of Exolysin in the mutant *Pch*Δ*exlBA* attenuated significantly the mortality of infected larvae (Figure 6B), suggesting the contribution of Exolysin in larvae death. We then set-up experiments to evaluate the toxin contribution in *D. melanogaster* killing by feeding the flies with high doses of bacteria. As a positive control, we used a well-known fly pathogen, *P. entomophila* [21, 32], and we included an isogenic mutant (*Pe*-*exlAmut*) with a plasmid insertion within *PSEEN2177* after residue 442 codon of ExlA^Pe^ _(1-1416)_. As previously reported [33, 34], female flies being fed with high doses of *P. entomophila* (OD_600_ = 100) died within 2 and 4 days. Interestingly, the *Pe-exlAmut* showed significantly lower killing capacity compared to the *P. entomophila* wild-type strain (Figure 7). When used in the same experimental conditions, *P. chlororaphis* PA23 provoked mortality with delay of several days compared to *P. entomophila* (Figure 7), and the contribution of ExlA was less pronounced, although still significant. Together, these results show that in two different *Pseudomonas* species the disruption of a gene encoding two-partner secretion toxin ExlA reduced bacterial toxicity toward two different insects showing the role of ExlA in both bacteria-insect pathogenic interactions.

**Figure 6.**
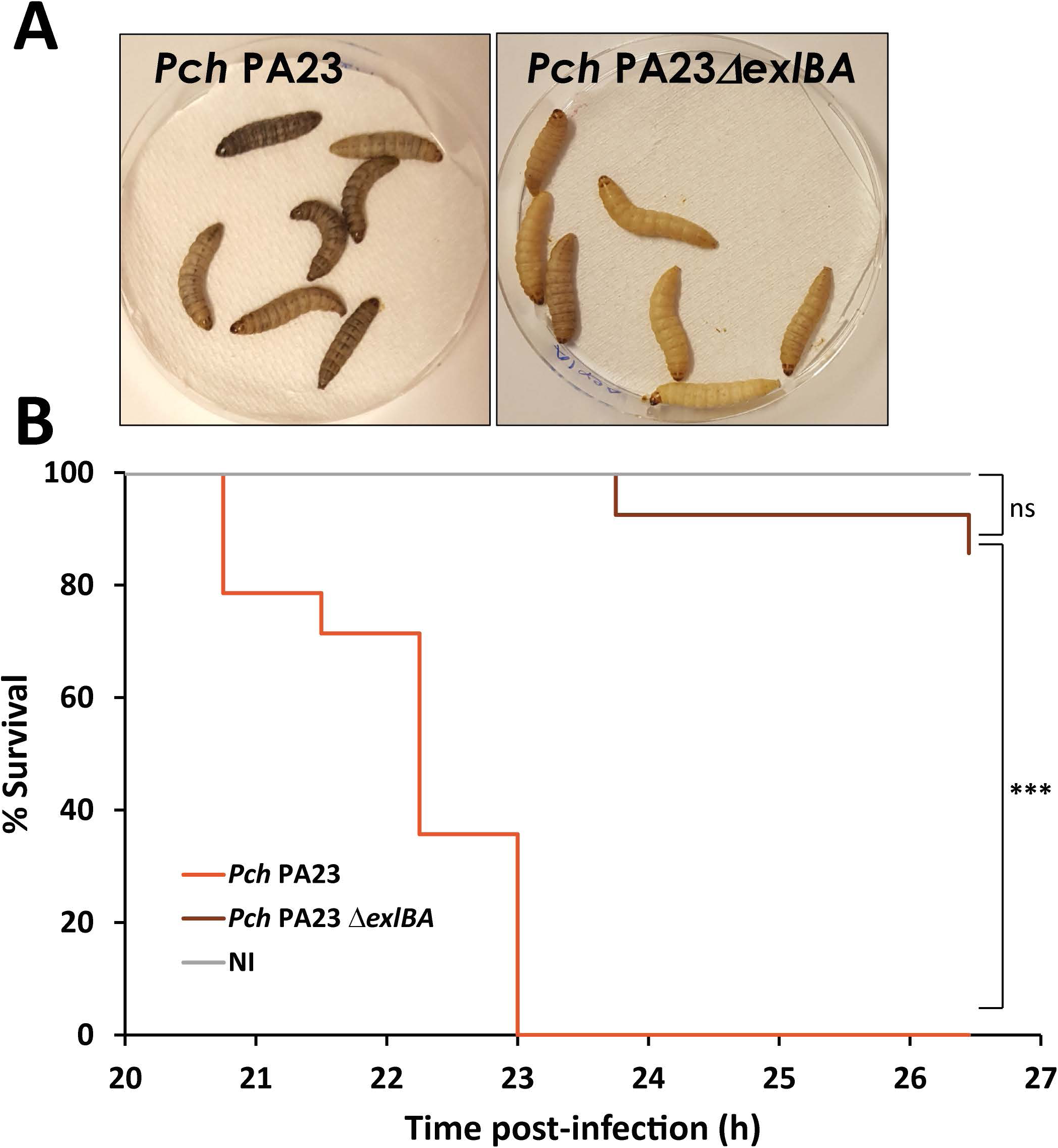
ExlA contributes to killing of wax moth larva *Galleria mellonella* by *P. chlororaphis* PA23. **A.** Representative experiment showing dead larvae after injection of *P. chlororaphis* PA23. Note the appearance of a dark color as indication of melanization. **B**. Survival rates of the larvae are expressed as percentage (%). The mortality was scored every 45 min. Larvae were injected with approx. 6×10^4^ bacteria in PBS, and let at 30 °C. Control larvae were injected with sterile PBS. Log rank test was used to determine statistically significant difference with P<0.001(***).

**Figure 7.**
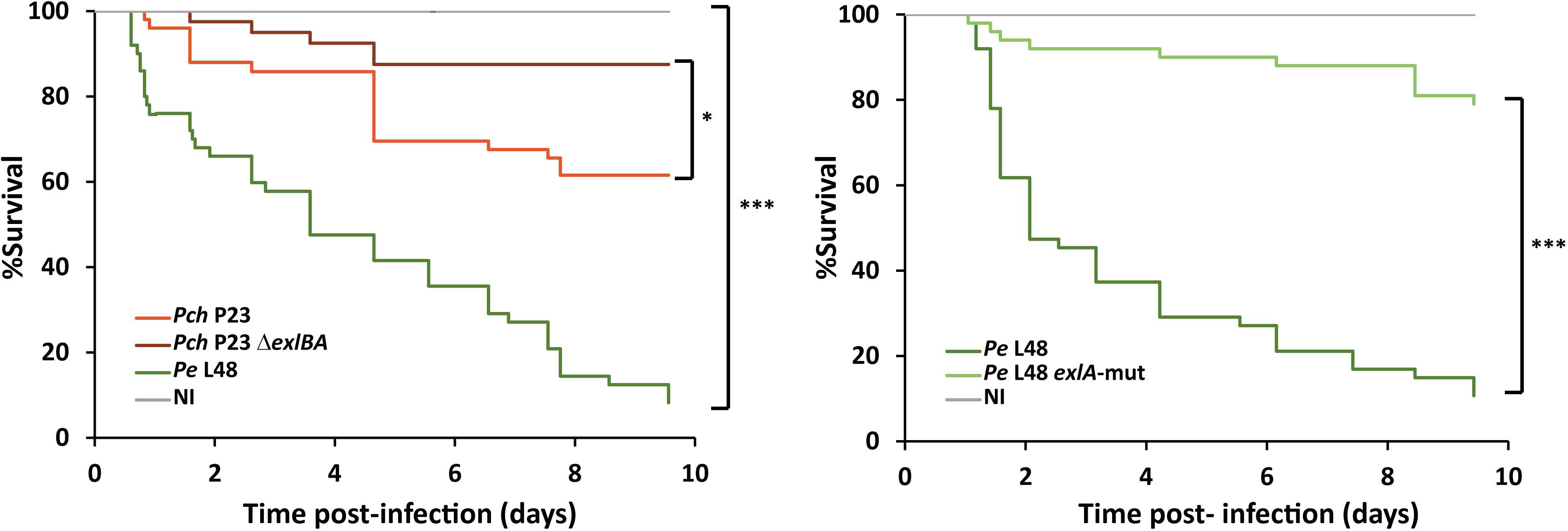
ExlA is important for *Drosophila melanogaster* oral infection by *Pseudomonas* sp. 4-8 days old female flies were fed with 1 % sucrose containing OD_600_ = 100 of indicated strains. In right panel infection with *P. chlororaphis* P23 (*Pch*) wild-type (in orange) or *exlBA* mutant (in brawn); in left panel flies infection with *P. entomophila* L48 (*Pe*) wild-type (dark green) or *exlBA* mutant (light green). Flies were incubated at 29 °C and survival was monitored up to 10 days. A solution of 1 % sucrose without bacteria was a control condition (grey line). Log rank test was used to determine statistically significant difference with P<0.05 (*) and P<0.001(***). P=0.0145 (between *Pch* and *Pch*Δ*exlBA*), P=0.0001 (between *Pe* and *Pe-exlBA* mut).

## Conclusions

In this work, we characterized ExlB-ExlA system, originally annotated as a TPS secreting a filamentous hemagglutinin adhesin, in a biocontrol bacteria *P. chlororaphis*. ExlA^*Pch*^ is responsible for *P. chlororaphis* cytotoxicity measured on human epithelial cells and mouse macrophages, with similar kinetics observed with ExlA-producing *P. aeruginosa*. The ExlA protein contributed to *P. chlororaphis* toxicity in *Galleria mellonella* larvae opening the possibility to modulate ExlA levels and to increase the use of *P. chlororaphis* in management of pest insects. The link of ExlB-ExlA system to known toxins (i.e. Fit toxin, or effectors of T6SS) and secondary metabolites known to be important in bacteria-insect interactions should be characterized. Finally, as the Vfr/cAMP pathway activates the *P. aeruginosa exlBA* expression, and the promoter of *P. chlororaphis exlB* displays no conserved Vfr DNA-binding site, the regulatory mechanisms governing *exlBA* operon expression in *P. chlororaphis* will be important to investigate. Finally, our work revealed that ExlA contributes to *P. entomophila* virulence toward *Drosophila* flies, which opens the question of ExlA activity in this bacteria and the relationship between ExlA and *P. entomophila* toxin Monalysin that displays pore-forming activity and provokes fly intestinal damage [33].

## Materials and Methods

### Bacterial strains, growth conditions and genetic constructions

*P. chlororaphis* PA23 [12] was a gift from Terasa de Kievit, University of Manitoba, Canada. *P. aeruginosa* IHMA87 is a urinary tract infection isolate and was previously shown to secrete Exolysin [19]. *P. entomophila* [32] was obtained from Bruno Lemaitre’s lab (EPFL, Switzerland) and the plasmid-insertion mutant in the *exlA^Pe^* gene homologue, PSEEN2177, was given by Isabelle Vallet-Galey (I2BC-CNRS, Paris, France). The *exlA^Pe^* mutant was verified by PCR and sequencing. For *P. chlororaphis* PA23 ΔexlBA (*RS20945/950*) mutant construct, a 900 bp fragment containing a frameshift fusion of the two genes was synthesized by Genewiz and then subcloned in *Eco*RI-*Hin*dIII of pEXG2, leading to pEXG2-Mut-Pchloro_*exlBA*. The plasmid was transferred into *P. chlororaphis* by triparental mating using pRK600 as a helper plasmid. For allelic exchange, cointegration events were first selected on LB plates containing rifampicin (25µg/ml) and gentamicin (25µg/ml) at 28 °C. Single colonies were then plated on NaCl-free LB agar plates containing 10 % (wt/vol) sucrose to select for the loss of plasmid. The sucrose-resistant strains were checked for gentamicin sensitivity and mutant genotype by PCR. The mutant strains harbor a deletion of 18 % of *exlA* and 32 % of *exlB* genes. Bacteria were grown in liquid LB medium (Becton Dickinson) at 28-30 °C (for *Pch* and *Pe*) or 37 °C (for *Pa*) with 300 rpm agitation. After overnight incubation, strains were diluted in LB medium to reach optical density measured at 600_nm_ (OD_600_) of 1.0 at 30 °C or 37 °C, respectively.

### Cytotoxicity tests

Epithelial lung carcinoma cell line A549 (ATCC CCL-185), A549-EGFP [35] and J774 macrophages were grown in DMEM or RPMI medium (Life Technologies) supplemented with 10 % fetal calf serum (Lonza). For cytotoxicity test, cells were seeded at 12,500 cells per well for A549 and 100,000 cells per well for J744 on black µclear 96-well plates (Greiner) and used 48 h later to obtain confluent monolayers. One hour before infection, medium was replaced by DMEM or RPMI (for J774 cells) without phenol red, supplemented with PI (Sigma, 1 µM). Cells were infected at a multiplicity of infection (MOI) of 10, unless otherwise specified. PI incorporation was followed by fluorescence measuring (excitation 544 nm/emission 590 nm) every 10 minutes with Fluoroskan Ascent FL2.5 Microplate Fluorometer (Thermo Corporation), during 10 hours at 30 °C. Data were presented as arbitrary fluorescence units (A.U.) as a function of time.

### Imaging of infection by confocal video microscopy

For confocal video microscopy experiments, cells were seeded at 100,000 cells per well on Lab-tek II 8-chambered coverslips (Dutscher Scientific) and used 48 h later to obtain confluent monolayers. One hour before infection, medium was replaced by DMEM without phenol red and supplemented with 1 µM PI. Cells were infected at a MOI of 10 and immediately observed by confocal video microscopy for 6 h at 30 °C. Video microscopy was performed on a spinning-disk inverted microscope (Nikon TI-E Eclipse) equipped with an Evolve EMCCD camera. The optical sectioning was performed by a Yokogawa motorized confocal head CSUX1-A1. Images were acquired using an illumination system from Roper Scientific (iLasPulsed) with a CFI Plan Fluor oil immersion objective (40X, N.A. 1.3). Z-series were generated every 5 min using a motorized Z-piezo stage (ASI) by acquiring 10 z-plane images with a step size of 1 μm. Microscope was controlled with MetaMorph software (Molecular Devices). Temperature, CO_2_, and humidity control was performed using a chamlide TC system (TC-A, Quorum technologies). Solid-state 491 and 561 nm lasers (iLas, Roper Scientific) and ET 525/50M (Chroma) and FF01-605/54 (Semrock) emission filters were used for excitation and emission of EGFP and PI fluorescence, respectively. The cells used for analysis were chosen at the initial step of recording. Analyses were performed by measuring the fluorescence intensity of each cell using the ROI Intensity Evolution plugin on ICY software.

### Plaque formation assay

*Acanthamoeba castellanii* (ATCC 1034) was cultured at 30 °C in flasks with 20 ml of PYG medium (peptone proteose 2 %, yeast extract 0.1 %, Na Citrate 2 H_2_O 0.1 %, CaCl_2_ 0.4 mM, MgSO_4_ 4 mM, Na_2_HPO_4_ 2.5 mM, KH_2_PO_4_ 2.5 mM, Fe(NH_4_)_2_(SO_4_)_2_ 0.05 mM Glucose 0.1 M). 100 μl of overnight bacterial culture were pelleted by centrifugation at 4000 rpm for 5 min and resuspended in 1 ml of M63 medium. 100 µl of this resuspension at OD_600nm_ of 0.1 were spread on M63 Glucose 0.2 % agar plates to form a bacterial lawn. The plates were dried 20 min. Amoeba’s cells were collected by centrifugation, washed once with M63 medium, and different number of amoebae cell in 5 µl M63 were deposited on the top of the agar plate. Plates were incubated at 30 °C for 5 days. The least number of *Acanthamoeba castellanii* cells deposited above that was able to form plaque on the bacterial lawns was defined as the minimum number of cells required for plaque formation in this study.

### Infections of *Galleria mellonella*

Infections of *Galleria* larvae were done as previously described [36] with some modifications. The bacterial dose of approx. 60000 bacteria/injection was evaluated in preliminary experiment as such to obtain larvae mortality within 40 h post-pricking. Incubations were done at 30 °C and larvae were counted every 45 min. The death was evaluated by the insusceptibility to touch by plastic tweezers. The dead larvae were removed from the dish. The experiment was repeated three times with 20 larvae used for each condition.

### *Drosophila* oral infections

*Drosophila melanogaster white* Canton-S (*w^CS10^*) flies were maintained at 25 °C. Oral infection was done on 4 to 8-day old female flies let starving in an empty vial for 3-4 h before being transferred to an infected vial containing bacterial solutions. *P. chlororaphis* PA23 (wild-type and the Δ*exlBA* mutant) and *P. entomophila* L48 (wild-type and *exlA* mutant) were grown in LB at 28-30 °C. Overnight bacterial cultures were pelleted and suspended into a sterile 1 % sucrose solution and adjusted to an OD_600_ of 100 corresponding to 6.6 × 10^10^ bacteria /ml. 200 µl of bacterial suspension was added to a filter paper disk on the top of a standard fly-feeding medium in the infected vials [37]. Flies were maintained at 29 °C and the survival was monitored over 2 weeks. Vials containing filter paper imbedded with 200 µl 1 % sucrose alone were used as negative control. Infection experiment was repeated three times with 40 - 50 flies per conditions tested, only one representative experiment is shown. The statistical analysis was performed with SigmaPlot software using a log rank statistic method.

### Lipid raft isolation

Epithelial cell line A549 was seeded in P100 flasks at 2 × 10^7^ cells/flask. Thirty minutes before infection, 7 ml of fresh DMEM media was added per plate, four plates were used per condition. Overnight bacterial cultures were diluted in LB medium and let grown up to OD_600_ of 1.1 before being added on A459 monolayer cells at an MOI of 10. Cells infected with *P. chlororaphis* PA23 wild-type and *exlBA* mutant were incubated at 30 °C in presence of 5 % CO_2_ for 3 h and 30 min. Cells infected with *P. aeruginosa* IHMA87 strain were incubated at 37 °C in presence of 5 % CO_2_ for 4 h. Infection was monitored by microscopy and stopped when approximately 70 % of the cells started to change the morphology and shrink. At the end of the infection, the culture medium was used for quantification of lactate dehydrogenase (LDH, Roche) to evaluate the level of cytotoxicity (cell permeabilization), while the infected A549 cells were washed twice with PBS and then resuspended into 1.6 ml of 50 mM HEPES, 150 mM NaCl, 5 mM EDTA, 1 % TritonX-100, pH7.4, containing Protease inhibitors cocktail (Complete, Roche). Resuspended A549 cells were incubated in ice for 1 h and kept in suspension by frequently vortexing. The debris were eliminated by low speed centrifugation at 1,000 g for 10 min at 4 °C and the supernatants were applied on a sucrose gradient. After ultracentrifugation at 40,000 rpm for 16 h at 4 °C, the lipid rafts are visible as a white cloudy band. Fractions of 1 ml were recovered from the top to the bottom of each tubes and loaded on a gradient 4-12 % SDS-PAGE Bis-Tris gel (BioRad). For Western Blot analysis, the proteins were transferred onto a PVDF membrane and incubated with mixture of rabbit polyclonal anti-ExlA antibodies. The mixture was composed by antibodies against ExlA-synthetic peptides diluted 500x [19], antibodies anti-ExlA-Cter and anti-ExlA-delta Cter both diluted 1 000x [25]. Mouse anti-Flotillin-2 antibodies (BD Biosciences) were used as lipid raft marker at a dilution of 1:10,000.

## Acknowledgments

We thank Marie-Odile Fauvarque and Emmanuel Taillebourg (IRIG, BGE, CEA Grenoble) for help with *Drosophila melanogaster* experiments, and Isabelle Vallet-Gely for providing the *P. entomophila* mutant. We would like to thank Emma Chastaing for her technical assistance for the experiments with amoebas and fungi during her master internship (MAP, Lyon). The work was supported by grants from Agence Nationale de la Recherche (ANR-15-CE11-0018-01), the Laboratory of Excellence GRAL, financed within the University Grenoble Alpes graduate school (Ecoles Universitaires de Recherche) CBH-EUR-GS (ANR-17-EURE-0003) and the Fondation pour la Recherche Médicale (Team FRM 2017, DEQ20170336705).

